# Srs2/PARI DNA helicase mediates abscission inhibition in response to chromatin bridges in yeast and human cells

**DOI:** 10.1101/2024.11.13.623343

**Authors:** Monica Dam, Nicola Brownlow, Audrey Furst, Coralie Spiegelhalter, Manuel Mendoza

## Abstract

The coordination of chromosome segregation with cytokinesis is crucial for maintaining genomic stability. Chromatin bridges, arising from DNA replication stress or catenated chromosomes, can interfere with this process, leading to genomic instability if not properly managed. Here, we uncover that the budding yeast DNA helicase Srs2 and its human homolog PARI delay the timing of abscission events in the presence of chromatin bridges. We demonstrate that Srs2 is essential for delaying abscission in yeast cells with chromatin bridges, and preventing their damage by cytokinesis. In human cells, PARI similarly plays a key role in delaying abscission events, such as midbody severing and actin patch disassembly during cytokinesis, in response to chromatin bridges caused by topoisomerase II inhibition. Our results also show that PARI functions within the Aurora B-mediated abscission checkpoint pathway. These findings reveal an evolutionarily conserved role of the Srs2/PARI DNA helicase in maintaining genomic integrity by modulating abscission timing in response to chromatin bridge formation.

## Introduction

The proper coordination between chromosome segregation and cell division (cytokinesis) is critical for maintaining genomic stability and supporting cell proliferation. Chromatin bridges that persist during cytokinesis can arise from various sources, including defects in DNA replication, chromatin condensation, DNA decatenation and by dicentric chromosomes formed by telomere fusion, among other anomalies (Ganem and Pellman, 2012; Mankouri et al., 2013). If these chromatin bridges are not adequately resolved, they can either be damaged during cytokinesis or cause cytokinesis failure, both of which have detrimental effects on cell proliferation.

Cytokinesis failure can result in binucleation and tetraploidy, conditions that are linked to the promotion of tumorigenesis (Fujiwara et al., 2005). Notably, a large fraction of human carcinomas are thought to originate from tetraploid cells, in part due to failed cytokinesis (Zack et al., 2013). Conversely, damage to unsegregated DNA during cytokinesis can lead to aneuploidy, gross chromosome rearrangements and impaired cell growth (Janssen et al., 2011). At the organismal level, these defects can contribute to severe pathologies such as T-cell lymphoma and microcephaly (Woodward et al., 2016; Martin et al., 2016). Therefore, chromatin bridges serve as both indicators of genomic instability and precursors to cellular transformation. Understanding how cells respond to chromatin bridges of different origins remains a critical question.

Given the potentially damaging consequences of chromatin bridges, cells have evolved mechanisms to mitigate their harmful effects. One such mechanism is the “NoCut” checkpoint, which delays the final step of cytokinesis, plasma membrane abscission, in budding yeast cells with chromatin bridges (Norden et al., 2006; Mendoza et al., 2009; Amaral et al., 2016). This checkpoint is conserved in animal cells, where it is known as the “abscission checkpoint” (Steigemann et al., 2009; Carlton et al., 2012; Agromayor and Martin-Serrano, 2013; Andrade and Echard, 2022). The NoCut-dependent delay in abscission is thought to protect chromatin bridges from damage and reduce the likelihood of abortive cytokinesis, which can occur if chromosomes obstruct the division site. Consequently, defects in the NoCut pathway in otherwise normal cells can lead to cytokinesis failure, tetraploidy, and genome rearrangements, all of which are hallmarks of cancer and may contribute to tumorigenesis.

In both yeast and animal cells, the function of the NoCut checkpoint depends on the association of the kinase Aurora-B with microtubules at the division site (the spindle midzone or intercellular bridge), suggesting that chromatin bridges are detected at this location. Aurora-B regulates the timing of abscission by controlling membrane remodelling events at the division site. In budding yeast, this regulation involves effectors such as Boi2, which influences the evolutionarily conserved exocyst complex that facilitates the fusion of secretory vesicles with the plasma membrane (Masgrau et al., 2017). In human cells, Aurora-B modulates abscission by regulating the endosomal sorting complex required for transport (ESCRT) III, a midbody-associated complex crucial for abscission timing in animal cells (Carlton et al., 2012; Addi et al., 2018). While the regulation of abscission timing downstream of Aurora-B is becoming clearer, the mechanisms by which chromatin bridges are sensed upstream of Aurora-B remain largely unknown.

Previous findings from our laboratory suggested that the yeast NoCut checkpoint does not merely respond to chromatin bridges, but specifically to bridges associated with DNA replication stress, possibly by detecting specific molecules associated with these bridges (Amaral et al., 2016, 2017). A similar phenomenon was also observed in human cells (Petsalaki et al., 2023). DNA helicases, known for their roles in DNA repair and replication stress resolution, have been implicated in resolving chromatin bridges in human cells, where they prevent bridge breakage and cytokinetic failure (Chan et al., 2018). Given these findings, we hypothesized that yeast helicases such as Srs2 could influence cytokinesis by resolving chromatin bridges during replication stress. Srs2 is a multifunctional helicase involved in DNA repair and homologous recombination regulation, particularly in dismantling Rad51 filaments (Marini and Krejci, 2010), and shares functional similarities with human helicases like PARI (Moldovan et al., 2012; Burkovics et al., 2016; Mochizuki et al., 2017). In this study, we explored the role of Srs2 in abscission and were initially expecting it to resolve chromatin bridges that would delay abscission. Surprisingly, we found that Srs2 has an additional function: it is required for the inhibition of abscission in response to chromatin bridges. This finding suggests that Srs2 plays an active role in the NoCut checkpoint, acting as a mediator of chromatin-based signaling during cytokinesis. Furthermore, we found that the human homolog PARI plays a similar role, delaying key events associated with abscission, including midbody severing and actin patch disassembly. These results suggest a conserved function for Srs2 and PARI in coupling chromatin bridge detection to the abscission machinery, extending our understanding of the molecular basis of NoCut signaling in both yeast and human cells.

## Results

### Srs2 promotes the resolution of anaphase chromatin bridges following replication stress

The budding yeast DNA helicase Srs2 is known for its role in inhibiting homologous recombination during S-phase and removing Replication Protein A (RPA) from chromatin (Pfander et al., 2005; Marini and Krejci, 2010; Dhingra et al., 2021), but its involvement in chromosome segregation and cytokinesis has not been explored. To investigate whether Srs2 contributes to the coordination of chromosome segregation and abscission, we monitored chromosome segregation and RPA localisation (visualised using histone H2B-mCherry and Rfa2-GFP) by time-lapse imaging in wild-type and Srs2-deficient budding yeast cells. Given the importance of Srs2 in suppressing replication intermediates during replication stress, we also examined chromosome segregation in *srs2Δ* cells exposed to hydroxyurea (HU), which perturbs DNA replication.

Treatment with a 3-hour pulse of HU generated late-segregating chromatin bridges in wild-type cells, as previously reported (Amaral et al., 2016), and this phenotype was exacerbated in cells lacking *SRS2* (Fig. **1A-B** and Fig. **S1**). Chromosome segregation was considered complete only when bridges were no longer detectable, while fragmented or discontinuous bridges were still classified as unresolved since stretched DNA could result in weak or absent nucleosome signals. In wild-type cells, 20% of log-phase cells entered anaphase (detected by nuclear elongation) with RPA foci, a percentage that increased to 40% under replication stress induced by HU and was further increased to 60% in the absence of Srs2 (Fig. **1C**). In wild type cells, RPA foci were primarily confined to the main nuclear masses; whereas in *srs2Δ* cells, over 20% of anaphase cells showed RPA foci located within chromatin bridges, regardless of prior HU exposure (**Fig. 1A** and **1D**). In cells with anaphase RPA foci, regardless of their subcellular location, chromatin bridges persisted longer than in cells without RPA foci, consistent with previous observations in unchallenged wild-type cells (Ivanova et al., 2020) (Fig. **1E**). The longest-lived bridges observed in HU-treated *srs2Δ* cells with anaphase RPA foci, indicating a severe delay in bridge resolution in the absence of Srs2. These findings suggest that the increased RPA on mitotic chromatin in *srs2Δ* cells, particularly following replication stress, may result from an accumulation of unresolved RPA-coated DNA intermediates formed during S phase. This could hinder the rapid resolution of chromatin bridges during anaphase.

**Fig. 1:**
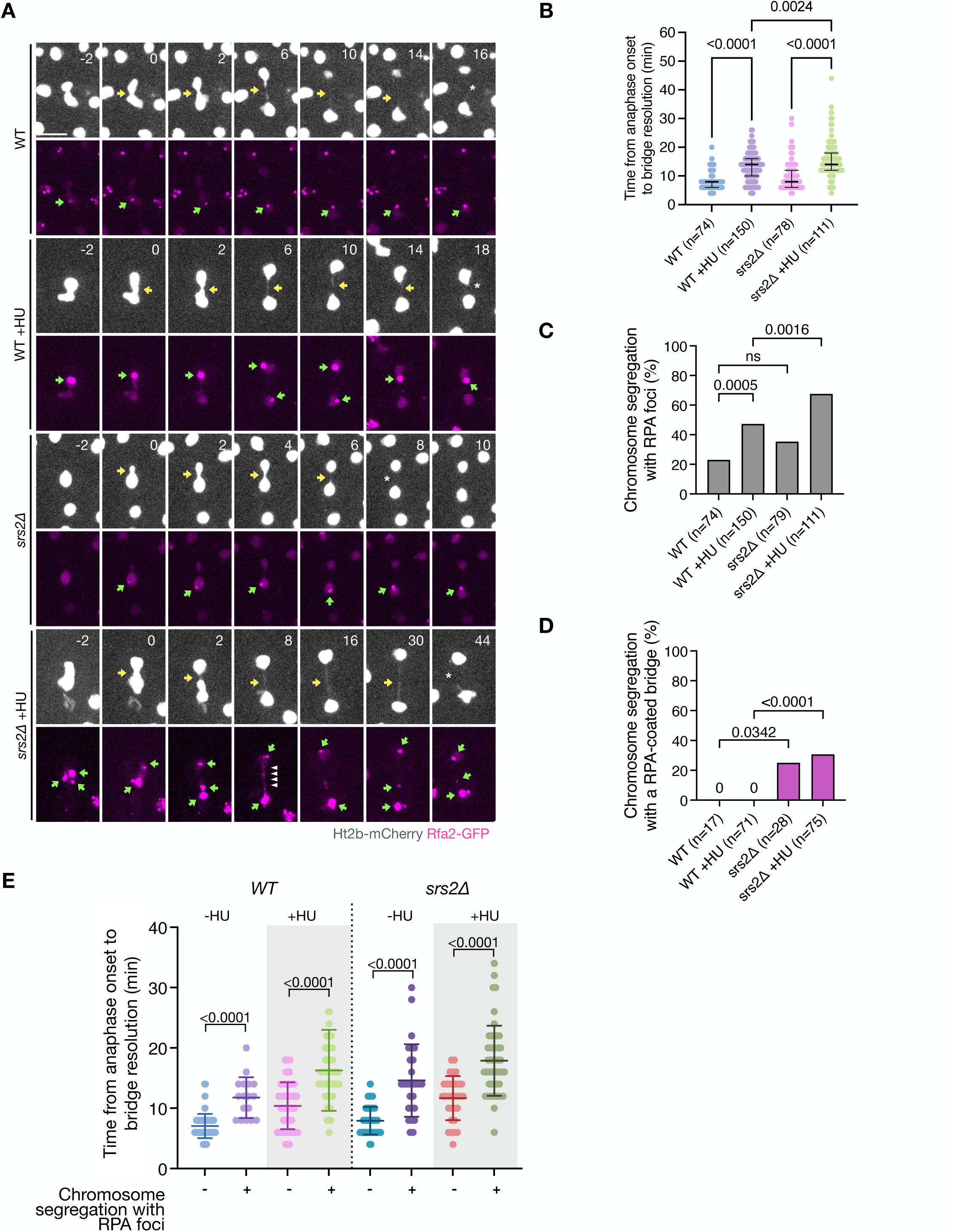
Srs2 promotes chromatin bridge resolution. (**A**) Chromosome segregation (Htb2-mCherry) and RPA foci formation (Rfa2-GFP) in cells of the indicated strains with and without previous exposure to HU. Selected frames are shown, and images are acquired every two minutes. Only cells with RPA foci are shown. Yellow arrows indicate chromatin bridges, green arrows point towards RPA foci, white arrowheads mark RPA bridges and asterisks note bridge resolution. Scale bar: 5 microns. (**B**) Time of bridge resolution (Htb2-mCherry) from the time of anaphase onset, defined by the rapid elongation of the nucleus. n =number of cells pooled from 2 independent experiments with similar results. P-values correspond to Dunn’s multiple comparison test. (**C**) Fraction of cells in (B) undergoing chromosome segregation with RPA foci. (**D**) Fraction of cells in (B) which segregated with RPA foci and had RPA-coated chromatin bridge. (**E**) Same data as in (B) but distinguishing between cells containing vs. not-containing RPA anaphase foci. P-values correspond to Mann-Whitney test.

### Srs2 is required for delaying abscission in the presence of chromatin bridges

To assess the impact of Srs2 loss in abscission timing, we used time-lapse confocal microscopy to monitor the ingression and resolution of the plasma membrane at the abscission site. For this, we used the fluorescent reporter GFP-CAAX, in which GFP is fused to the membrane-targeting CAAX motif of Ras2. The spindle pole protein Spc42 fused to GFP was used to monitor spindle elongation, which marks the start of anaphase. As previously described (Amaral et al., 2016), the morphology of membranes labelled with GFP-CAAX allows visualisation of abscission progression: following anaphase onset, the actomyosin ring begins to constrict the bud neck, leading to membrane ingression and eventually separation. The separation of the mother and daughter membranes marks abscission (Fig. **2A**). The relative timing of late mitotic events, including anaphase onset, nuclear division, membrane ingression and abscission, is summarised in Fig. **S2A**.

**Fig. 2:**
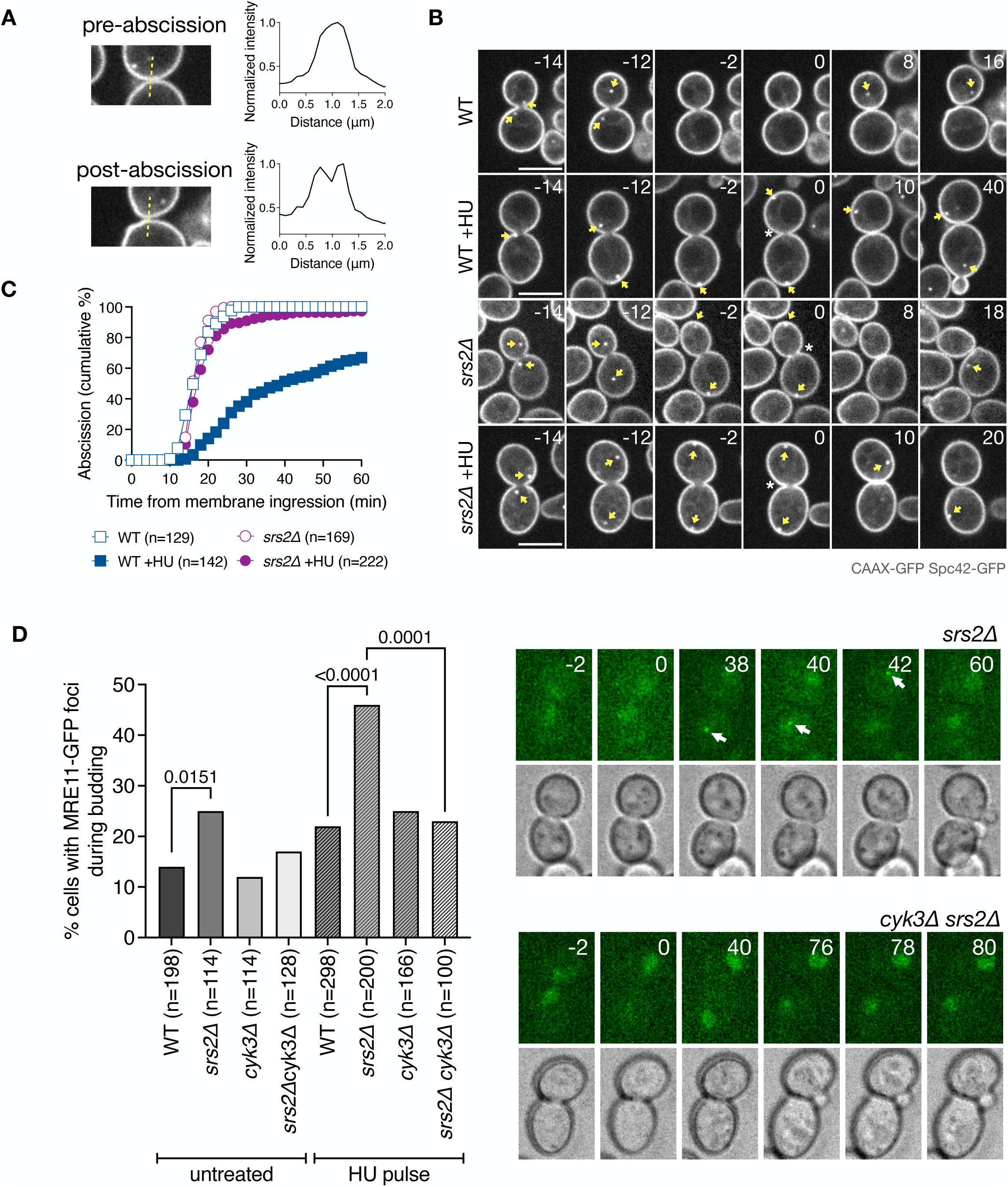
Srs2 prevents abscission-dependent DNA damage following replication stress. (**A**) Abscission is scored by measuring the distribution of GFP-CAAX intensity values across the mother-daughter cell axis at the bud neck. Upper panel: cells that have yet to abscise show a single peak of intensity at the bud neck. Lower panel: membrane separation is marked by two distinct peaks in GFP intensity. (**B**) Membrane ingression and abscission in WT and *srs2Δ* cells with or without an HU pulse. Cells express the plasma membrane marker CAAX-GFP and the spindle pole marker Spc42-GFP (yellow arrows). Time is indicated in minutes; 0 min indicates the start of membrane ingression. Asterisk specifies membrane ingression. White arrowhead marks complete constriction of the plasma membrane (abscission). Single Z-slices are shown, but 12 Z-slices spaced 0.3 μm, spanning the whole cell were used for image analysis. Scale bar 5 μm. (**C**) Fraction of cells that complete abscission from the time of membrane ingression. n =number of cells pooled from N =3 independent experiments with similar results. WT vs. WT +HU, and WT +HU vs. *srs2Δ* +HU, P <0.0001, Mann-Whitney test. (**D**) Frequency of Mre11–GFP focus formation after cytokinesis in the indicated strains and conditions. P-values correspond to Fisher’s exact test. Representative images of cells expressing Mre11–GFP, after cytokinesis following a HU pulse. The arrow points to a nuclear Mre11 focus.

In unchallenged WT cells, chromatin bridges resolve before membrane ingression, whereas in HU-treated cells, bridges resolve around the time of cytokinesis, coinciding with membrane ingression (Fig. **S2B**). In wild-type cells, the median time from membrane ingression to abscission is 16 min, but this is significantly delayed to 24 min upon HU treatment (Fig. **2B-C**). This aligns with our previous findings indicating that DNA replication stress delays bridge resolution and abscission through the NoCut checkpoint (Amaral et al., 2016). However, in the presence of HU-induced chromatin bridges, *srs2Δ* cells exhibited accelerated membrane ingression and failed to delay abscission (Fig. **2B-C** and Fig. **S2B**). These findings indicate that Srs2 is crucial for inhibiting cytokinetic abscission under conditions of DNA replication stress.

The DNA double strand break repair protein Mre11 forms nuclear foci indicative of DNA damage. In NoCut-deficient cells with chromatin bridges, Mre11-GFP foci accumulate after cytokinesis, likely due to chromatin bridge breakage (Amaral et al., 2016). To determine whether the lack of coordination between chromosome segregation and abscission in Srs2-deficient cells results in DNA damage after cytokinesis, we visualised the formation of DNA damage foci using Mre11-GFP. In normally cycling cells, approximately 14% of wild-type cells exhibit Mre11 foci after nuclear division, with this fraction increasing to 25% in the absence of *SRS2*. To test if increased damage in *srs2Δ* cells is due to replication stress-induced lesions in the main nucleus or to cytokinesis-induced damage of chromatin bridges, abscission was delayed by deletion of the cytokinesis gene *CYK3*, which delays septum formation and subsequent abscission (Amaral et al., 2016; Onishi et al., 2013). Notably, the fraction of *srs2Δ* cells with DNA damage was reduced to near wild-type levels in *srs2 cyk3* cells (Fig. **2D**, “untreated”). This suggests that premature cytokinesis causes damage to chromatin bridges in Srs2-deficient cells. Following replication stress induced by HU, the fraction of wild-type cells exhibiting Mre11 foci was increased from 24% in wild type to 40% in *srs2Δ* cells; also in this case, delayed abscission in *CYK3*-deficient cells was able to reduce the fraction of cells with Mre11 foci (Fig. **2D**, “HU pulse”). These results demonstrate that Srs2 is required to delay cytokinesis and prevent chromatin bridge damage in response to replication stress.

Next, we investigated if Srs2 is involved in the abscission delay caused by catenated chromatin bridges by inactivating topoisomerase II using the temperature-sensitive allele *top2-4*. These mutants display chromatin bridges in 100% of anaphases, and delay abscission is a manner dependent on Aurora-B (Amaral et al., 2016). Cells were grown at 25°C and synchronised in G1 with α-factor for 2 hours. G1 arrested cells were then released into pre-heated media at 37°C and placed in a pre-heated chamber at 37°C for imaging. Wild-type and *srs2Δ* cells expressing the GFP-CAAX plasma membrane marker exhibited similar abscission timing (Fig. 3A-B). Consistent with previous findings (Amaral et al. 2016), inactivation of *top2-4* resulted in significant impairment in membrane resolution, with 92% of cells failing to complete abscission within 60 minutes. Remarkably, this impairment was partially but significantly rescued in cells lacking Srs2, reducing the failure rate from 92% to 41% (Fig. **3A-B**). These findings suggest that Srs2 plays a critical role in inhibiting abscission in cells challenged with both HU-induced and catenated chromatin bridges.

**Fig. 3:**
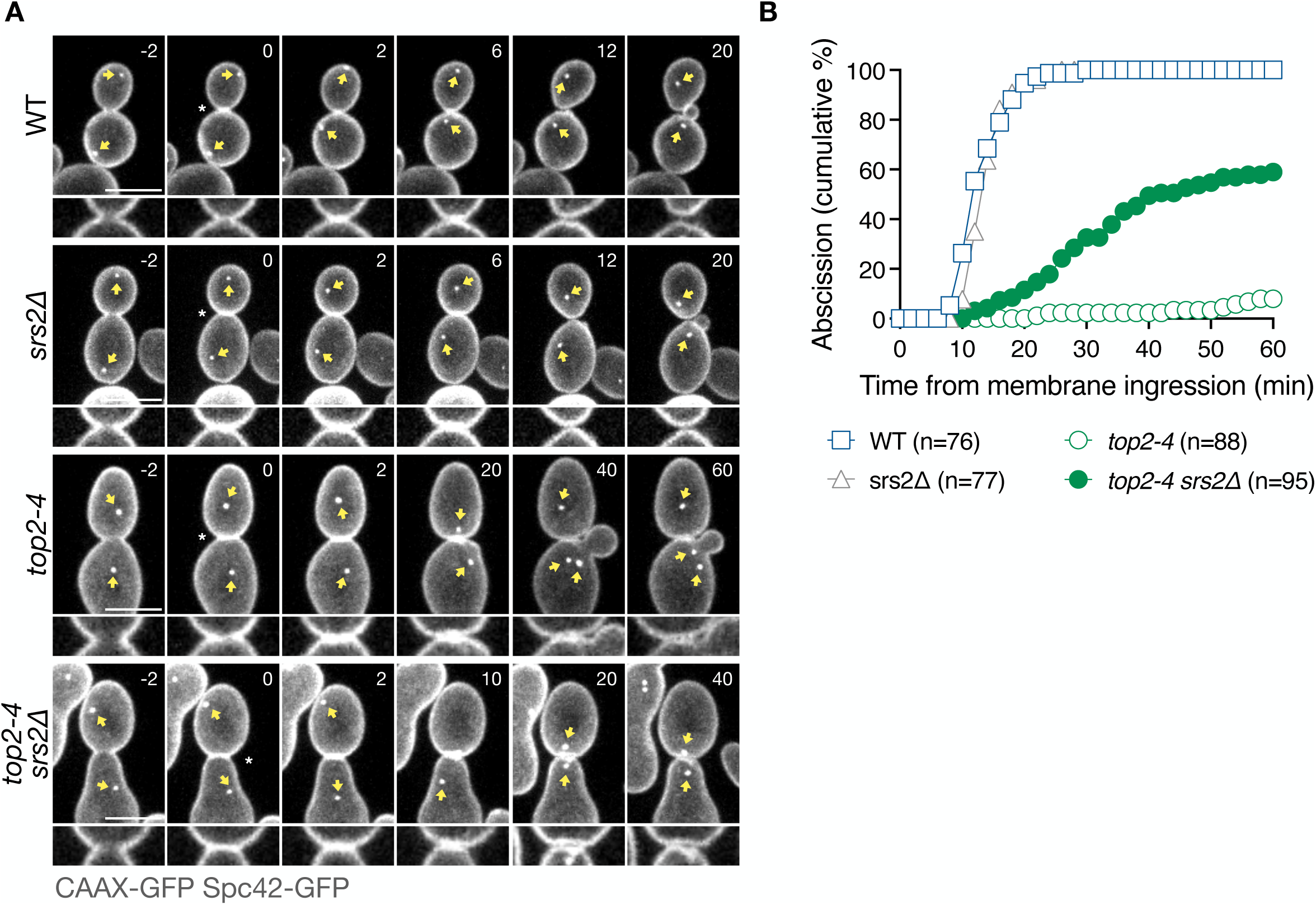
Srs2 promotes abscission inhibition in Topoisomerase II-deficient cells. (**A**) Membrane ingression and abscission in cells of the indicated genotypes. Time in minutes, 0 min indicates the start of membrane ingression. Yellow arrow marks the position of spindle poles. Asterisk specifies membrane ingression. White arrowhead marks complete separation of the plasma membrane. Time is indicated in minutes; 0 min indicates the start of membrane ingression. Asterisk specifies membrane ingression. White arrowhead marks complete constriction of the plasma membrane (abscission). Single Z-slices are shown, but 12 Z-slices spaced 0.3 μm, spanning the whole cell were used for image analysis. Lower Zoom-in panel shows the dynamics of membrane ingression at the central Z-slice. Scale bar 5 μm. (**B**) Fraction of cells that complete abscission from the time of membrane ingression. n =cells pooled from N =3 independent experiments with similar results. WT vs. *top2-4*, and *top2-4* vs. *top2-4 srs2Δ*, P <0.0001, Mann-Whitney test.

### Association of Srs2 with PCNA is required for full abscission inhibition

Srs2 is recruited to the replication fork during S-phase by SUMOylated PCNA, through its C-terminal SUMO-interacting motif (SIM) and PIP-box (Armstrong et al., 2012; Kolesar et al., 2012). To investigate whether this interaction is essential for the role of Srs2 in NoCut function, we first deleted the SIM of Srs2 and assessed abscission timing. Interestingly, deletion of the SIM alone was not sufficient to rescue abscission failure in *top2-4* mutants (Fig. **4A**). Although SIM-defective Srs2 has a lower affinity to SUMOylated PCNA, it can still interact with non-SUMOylated PCNA (Armstrong et al., 2012). We then deleted both the PIP-box and the SIM, thereby disrupting both interactions with SUMO and PCNA. The absence of both the SIM and PIP-box partially but significantly rescued the fraction of cells able to complete abscission, increasing it to 28% (Fig. **4A**). These findings suggest that interacting with PCNA is critical for Srs2 to fully promote a NoCut-mediated abscission delay in the presence of catenated bridges.

**Fig. 4:**
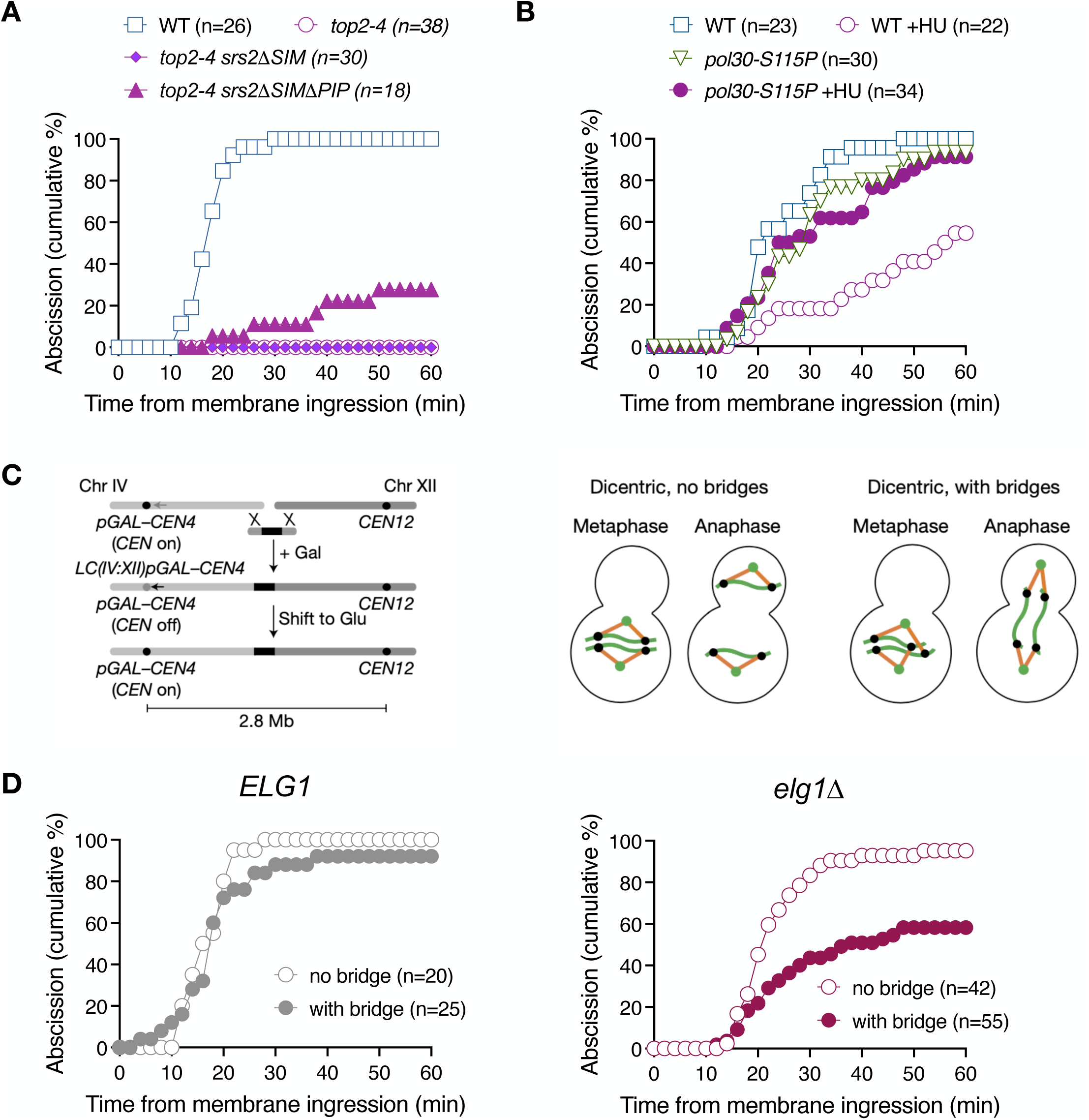
Chromatin association of Srs2 and PCNA promote inhibition of abscission in response to chromatin bridges. (**A**) Abscission dynamics of *srs2ΔSIM* and *srs2ΔSIMΔPIP* in the presence of catenated bridges. WT vs. *top2-4*, <0.0001; *top2-4* vs. *top2-4 srs2ΔSIM,* P =0.2990; *top2-4* vs. *top2-4 srs2ΔSIMΔPIP*, <0.0001, Mann-Whitney test. (**B**) Inactivation of cold-sensitive PCNA (*pol30-S115*) in cells exposed to DNA replication stress. WT+HU vs. *pol30-S11P*+HU, P =0.0004, Mann-Whitney test. (**C**) Schematic describing how the conditionally dicentric strain was generated, and how dicentric bridges are formed due to biorientation of the kinetochores. Large-budded cell in metaphase and anaphase, green dot – spindle pole, black dot – kinetochore, orange line – kinetochore microtubule, green line – dicentric chromosome. (**D**) Abscission dynamics in dicentric strains of the indicated genotypes. Dicentric bridges were imaged with Htb2-mCherry and abscission was scored with GFP-CAAX. *ELG1* cells with bridge vs. *elg1Δ* with bridge, P <0.0001, Mann-Whitney test. n= number of cells pooled from one (A, B), two (D, *ELG1*) or three (D, *elg1Δ*) independent time-lapse experiments.

To determine if PCNA is required during cytokinesis to modulate abscission timing, we used a cold-sensitive mutant of PCNA (*pol30-S115P*). Due to weakened interactions within the Pol30 homotrimer, this mutant shows reduced chromatin accumulation and does not grow at 14 °C (Ayyagari et al., 1995; Johnson et al., 2016). Wild-type and *pol30-S115P* cells were grown at the permissive *pol30-S115P* temperature of 30°C and exposed to HU for two hours, before shifting to the restrictive temperature of 14°C. Following HU exposure, *pol30* mutants were significantly less delayed in abscission than wild-type cells (Fig. **4B**). Pol30 inactivation in otherwise unchallenged cells (without HU) did not affect abscission timing. Together, these results further suggest that PCNA integrity, and likely its association with Srs2 on chromatin, is important for the NoCut checkpoint.

We next tested whether association of PCNA with chromatin is not only necessary but also sufficient to inhibit abscission. We took advantage of a conditional dicentric chromosome, generated by end-to-end fusion of chromosomes IV and XII, in which centromere 4 is kept inactive thanks to activation of the strong *GAL1,10* promoter. On activation of *CEN4* in glucose-containing media, chromatin bridges (visualised with Htb2-mCherry) form in 50% of anaphases (Fig. **4C**). These bridges do not trigger a NoCut response, possibly due to their failure to recruit a NoCut-activating factor (Amaral et al., 2016). We asked whether retaining PCNA and Srs2 on chromatin could render dicentric chromatin bridges detectable by the NoCut checkpoint. After replication termination, PCNA is unloaded from chromatin by the Replication Factor C-like complex Elg1. In cells lacking Elg1, PCNA remains associated with chromatin (Kubota et al., 2013).

We monitored abscission in conditionally dicentric mutants with wild-type *ELG1* and confirmed that abscission proceeds with near-wild-type efficiency, regardless of the presence of dicentric bridge (Fig. **4D**, *ELG1*). Similarly, *elg1Δ* cells without chromatin bridges completed abscission with dynamics comparable to wild-type cells (Fig. **4D**, *elg1Δ,* “no bridge”), indicating that PCNA retention on chromatin alone does not affect abscission timing. However, in the presence of a dicentric bridge, *elg1Δ* mutants exhibited delayed abscission, with approximately 50% of cells unable to resolve the membrane within 60 minutes (Fig. **4D**, *elg1Δ,* “with bridge”). This finding suggests that PCNA retention, or the persistence of PCNA-associated factors, specifically on dicentric chromatin bridges is necessary to inhibit abscission, rather than PCNA retention on chromatin in general.

### The Srs2 human homolog PARI delays midbody severing in response to chromatin bridges

Like Srs2, the human protein PARI acts as an anti-recombinase and prevents the accumulation of recombination intermediates that could lead to genomic instability during DNA replication (Burkovics et al., 2016; Moldovan et al., 2012; Mochizuki et al., 2017). To investigate the role of PARI in the human NoCut / abscission checkpoint, we developed an assay to measure chromatin bridge formation and abscission in HeLa cells. Cells were synchronised in the cell cycle using the double thymidine method. Cells were treated with thymidine for 16 hours to inhibit deoxynucleotide synthesis, causing arrest in S phase. This was followed by an 8-hours release into normal culture media and a subsequent 16-hour thymidine treatment to achieve a more synchronised block at the G1/S boundary. Following release from the double thymidine block, cells progressed to G2/mitosis after approximately 7.5 hours and entered cytokinesis 4.5 hours later (Fig. **5A**). To induce chromatin bridges with minimal S-phase perturbation, a low dose of the catalytic topoisomerase II inhibitor ICRF-193, which causes both catenated chromatin bridges and ultrafine (UFBs) (Germann et al., 2013; Bhowmick et al., 2019), was added during late G2 phase (Fig. **5A**).

**Fig. 5:**
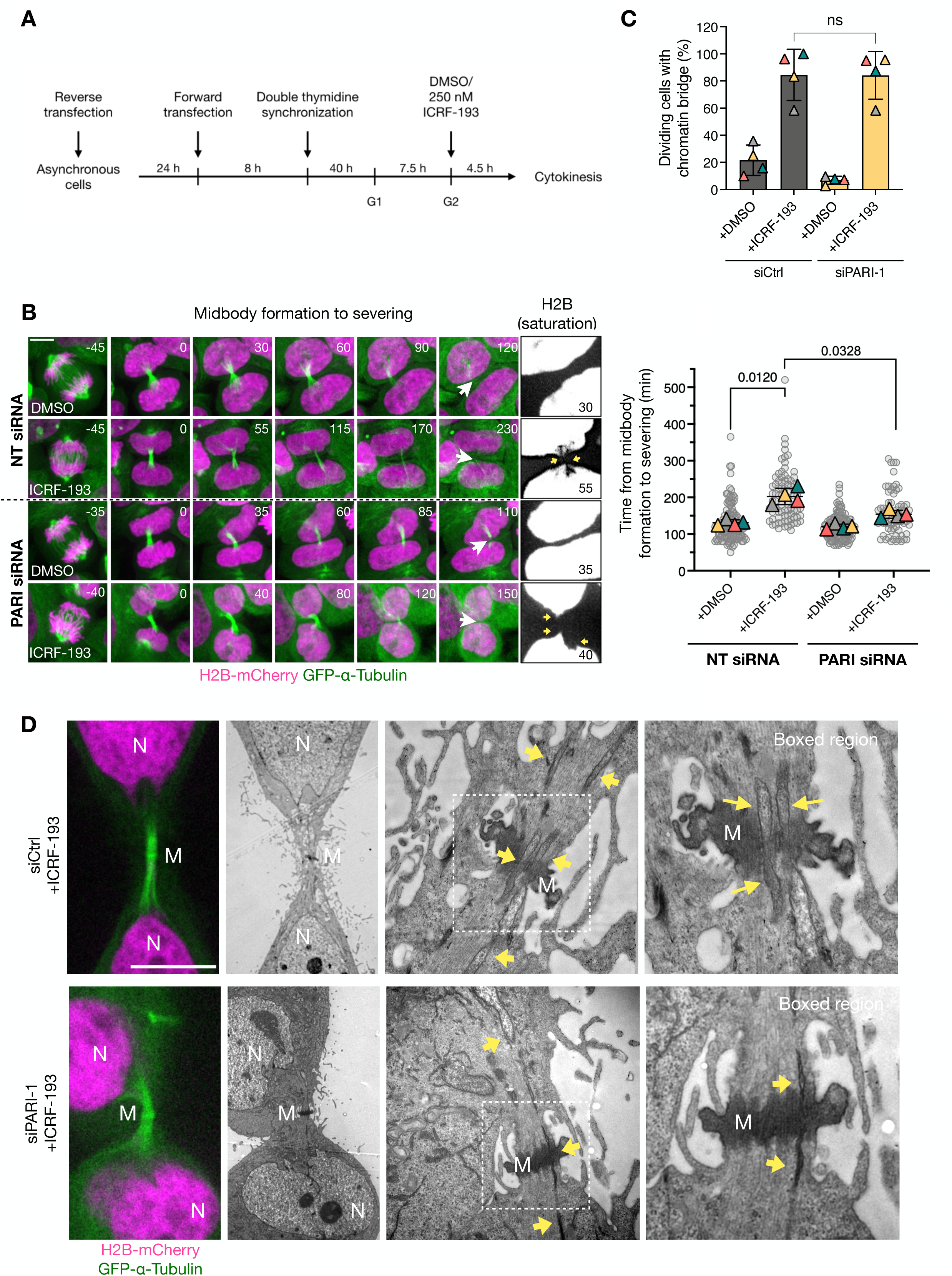
PARI is required to delay midbody severing in cells with catenated bridges. (**A**) Experimental workflow to synchronise siRNA-transfected cells and inducing catenated chromatin bridges during cytokinesis by the addition of ICRF-193 during G2. (**B**) Percentage of dividing cells containing chromatin bridges (H2B-mCherry) under control (siCtrl) or PARI depletion (siPARI-1), treated with either DMSO or ICRF-193. (**C**) Time-lapse images of cells expressing H2B-mCherry (magenta) and GFP-α-Tubulin (green) transfected with control or siPARI and treated with either DMSO or ICRF-193. High-contrast, saturation-adjusted projections of H2B-mCherry fluorescence images are shown to enhance the visualization of chromatin bridges. Arrowheads mark midbody severing. Scale bar, 5 um. The SuperPlot shows the quantification of the midbody lifetime (time from midbody formation to severing). Each cell is represented as a grey dot, the mean of each independent experiment is represented as a triangle. Student’s paired t test (mean ±SD, *P <0.05, **P <0.01; ***P <0.001; ****P <0.0001, N =4). (**D**) ICRF-193-treated cells stably expressing H2B-mCherry and GFP-alpha-Tubulin, transfected with control or PARI-1-specific siRNAs. Yellow arrows point towards chromatin, and boxed regions zoom in on the Flemming body. N = nucleus, M = midbody.

PARI was depleted in HeLa cells using siRNA. Since neither commercial nor in-house polyclonal antibodies detected the PARI protein at the expected 65 kDa size, RT-qPCR confirmed an 80% reduction in PARI mRNA levels compared to cells transfected with control siRNA (Supplementary Fig. **S3**).

Next, we evaluated abscission timing by measuring midbody stability. The duration from midbody assembly (marked by the constriction of central spindle microtubules into a bundle), to midbody severing is known to increase in the presence of chromatin bridges and serves as a proxy for abscission timing (Steigemann et al., 2009; Carlton et al., 2012; Advedissian et al., 2024). We term this duration “midbody lifetime”. To assess midbody lifetime, we conducted time-lapse spinning disk microscopy on synchronised HeLa cells stably expressing GFP-α-tubulin. Chromatin was visualised using H2B-mCherry.

Our results demonstrated that in untreated HeLa cells, the average midbody lifetime is 130 minutes. However, in the presence of ICRF-193, midbody severing is significantly delayed to approximately 200 minutes. Remarkably, in the absence of PARI, this delayed midbody severing is partially but significantly reduced, to around 150 minutes (Fig. **5B**). The severing of midbody microtubules in PARI-depleted cells occurred in the presence of chromatin bridges, as determined by H2B-Cherry ( Fig. **5B-C**) and confirmed using correlative light and electron microscopy (CLEM) (Fig. **5D** and Supplementary Fig. **S4**). Continuous chromatin was observed traversing the intercellular bridge (ICB), passing through the microtubule-dense midbody and Flemming body in cells treated with ICRF-193. The microtubules of the midbody remained straight and ordered along the ICB, even in the presence of a chromatin bridge. Furthermore, depletion of PARI did not appear to disrupt the structure of the ICB during cytokinesis, with morphologies similar to those observed in cells with intact PARI (Fig. **5D**).

To determine whether the role of PARI in regulating midbody lifetime is specific and not due to off-target effects of the siRNA used, we designed a siPARI-1-resistant version of PARI. The putative seed region of siPARI-1 was identified, and three silent mutations were introduced that did not alter the wild-type protein sequence but rendered the mRNA resistant to degradation by siPARI-1 (Supplementary Fig. **S5A-B**). Additionally, an eGFP tag was added to the N-terminus of PARI. We generated HeLa Kyoto cell lines stably expressing either eGFP or siPARI-1-resistant eGFP-PARI (eGFP-PARI^R^) (see Methods). These cells were depleted of endogenous PARI using siPARI-1 and treated with ICRF-193 as described. The midbody and microtubules were visualised using SiR-Tubulin dye, and midbody lifetime was measured from assembly to severing. In cells expressing eGFP, siPARI-1 treatment significantly reduced midbody lifetime, consistent with previous observations in control HeLa cells. However, in cells expressing eGFP-PARI^R^, siPARI-1 did not reduce midbody lifetime (Supplementary Fig. **S5C-D**). These results suggest that the human homolog of Srs2, PARI, specifically delays midbody severing in the presence of chromatin bridges induced by topoisomerase II inactivation.

### PARI does not stabilise the midbody upon NPC defects

It has been shown that defects in nuclear pore assembly, such as those caused by depleting the basket component NUP153, can delay midbody severing without the presence of chromatin bridges (Mackay et al., 2010). To determine whether PARI is involved in this response, we partially co-depleted NUP153 along with PARI in HeLa cells using siRNA (see Methods). The NUP153 protein level was assessed by Western blotting and was found to be partially depleted after transfection; co-depletion with PARI did not alter these levels (Supplementary Fig. **S5B**). Cells were synchronised in G1/S with thymidine for 24 hours and then released for 16 hours. NUP153-depleted cells distinctively arrested in cytokinesis, exhibiting cytoplasmic “abscission checkpoint bodies” (ACBs), which are indicative of an active abscission checkpoint (Strohacker et al., 2021) in cells treated with both control and PARI-targeting siRNAs (Supplementary Fig. **S5A, C**). These results suggest that PARI is not required for midbody stabilisation in response to nuclear pore assembly defects, but rather plays a specific role in delaying midbody severing in the presence of chromatin bridges.

### PARI promotes actin stabilisation at the intercellular bridge

In addition to delaying midbody severing, F-actin has been shown to promote abscission checkpoint-mediated delay by accumulating at the intercellular bridge (ICB) as actin patches, which stabilise the canal in the presence of chromatin bridges (Steigemann et al., 2009; Dandoulaki et al., 2018; Bai et al., 2020). To determine whether PARI regulates this process, we examined F-actin dynamics in HeLa cells stably expressing H2B-mCherry and GFP-actin.

In control cells (siCtrl + DMSO), actin accumulated at the cleavage furrow during anaphase and gradually dispersed as cytokinesis progressed. Depletion of PARI (siPARI-1 + DMSO) did not significantly alter actin behavior, as actin cleared from the division site with similar kinetics as in control cells (Fig. **6A-B**, **S7**, and Supplementary Movies **S1** and **S2**). In contrast, treatment with ICRF-193 (siCtrl + ICRF-193) led to a prolonged retention of actin at the division site, in patches often associated with H2B-labeled chromatin, consistent with delayed cytokinesis and the presence of chromatin bridges. These actin patches persisted for at least 200 minutes after membrane ingression, and even 700 minutes post-ingression, only 25% ICR-193-treated cells had disassembled them (Fig. **6A-B**, and Supplementary Movie **S3**). Interestingly, in PARI-knockdown cells, actin patches often formed but failed to persist in most cells, with 60% of cells lacking actin patches 700 minutes after membrane ingression (Fig. **6A-B**, and Supplementary Movie **S4**). These findings indicate that PARI promotes the formation of actin patches at the ICB, facilitating actin-mediated stabilization of the canal in the presence of chromatin bridges.

**Fig. 6:**
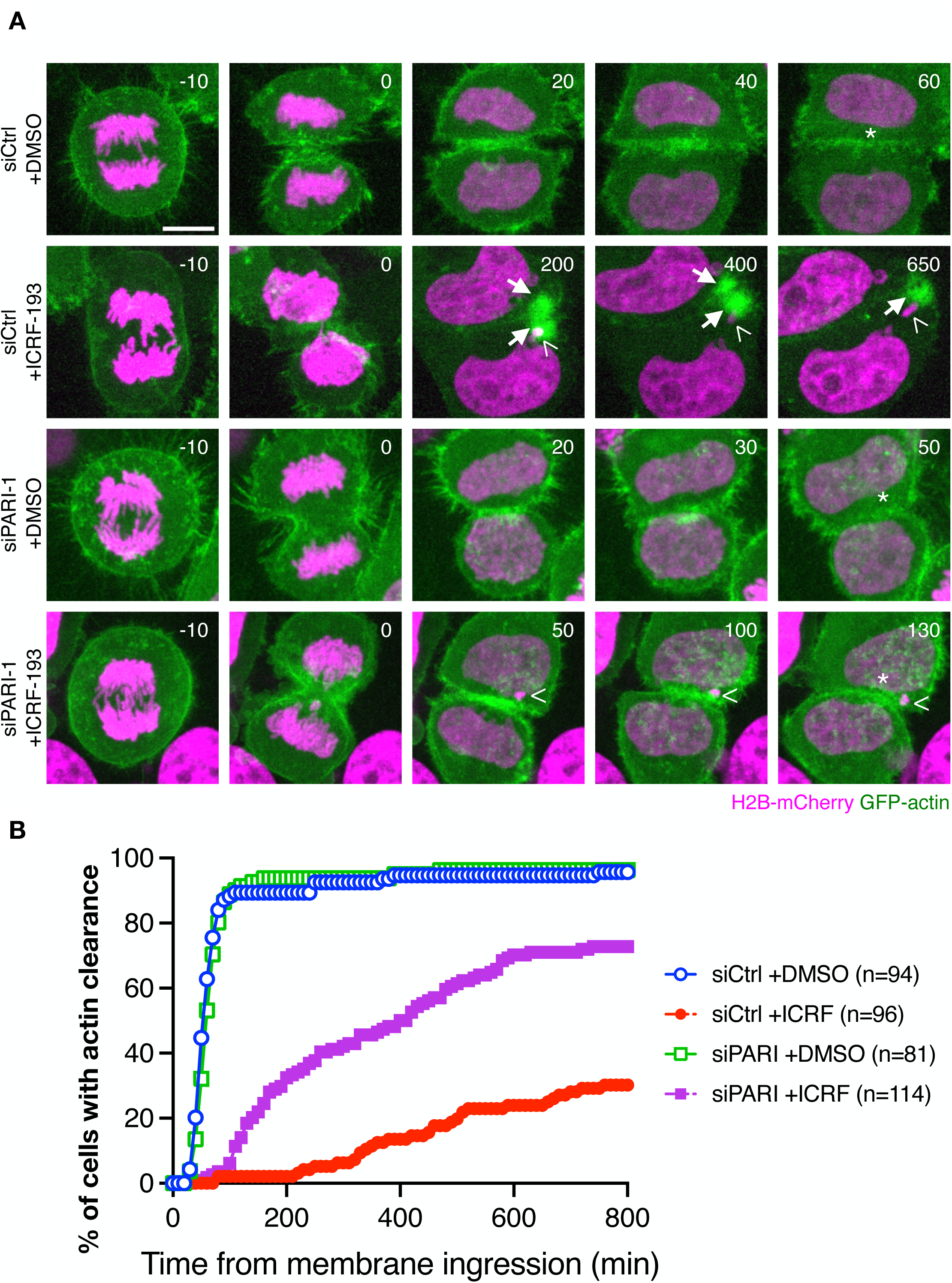
PARI promotes actin-patch clearance at the intracellular bridge in cells with chromatin bridges. (**A**) HeLa cells progressing from anaphase to membrane ingression and actin-patch disassembly (GFP-actin) with or without the presence of a catenated chromatin bridge (H2B-mCherry) induced by ICRF-193. Arrows point to actin patches, and arrowheads point to chromatin bridges. (**B**) Quantification of actin clearance from the division plane. n =cells pooled from N =3 independent experiments with similar results. siCtrl +DMSO vs. siCtrl +ICRF-193, P <0.0001; siCtrl +ICRF-193 vs. siPARI-1 +ICRF-193, P <0.0001, Mann-Whitney test.

### PARI regulates midbody stability through an Aurora B-dependent pathway

To determine whether catenated bridges induce a delay in midbody severing dependent on the Aurora B-mediated abscission checkpoint, and to investigate if PARI is a component of this checkpoint, we inhibited Aurora B at the time of cytokinesis. HeLa cells stably expressing H2B-mCherry and GFP-α-Tubulin were synchronised using the double thymidine block and treated with ICRF-193 following the standard protocol. Based on previous observations, midbody severing typically occurred approximately 120 minutes after the start of acquisition. At this point, the Aurora-B inhibitor hesperadin was added (Fig. **7A-B**). As expected, control cells treated only with ICRF-193 exhibited a mean midbody lifetime of 200 minutes, and Aurora B inhibition accelerated midbody disassembly, reducing the mean midbody lifetime to 90 minutes. In the absence of PARI, the midbody lifetime was similarly reduced, consistent with our previous observations in the presence of catenated bridges. Notably, co-depletion of PARI and inhibition of Aurora B did not further reduce the already shortened midbody lifetime, suggesting that Aurora B and PARI function within the same pathway to regulate midbody disassembly.

**Fig. 7:**
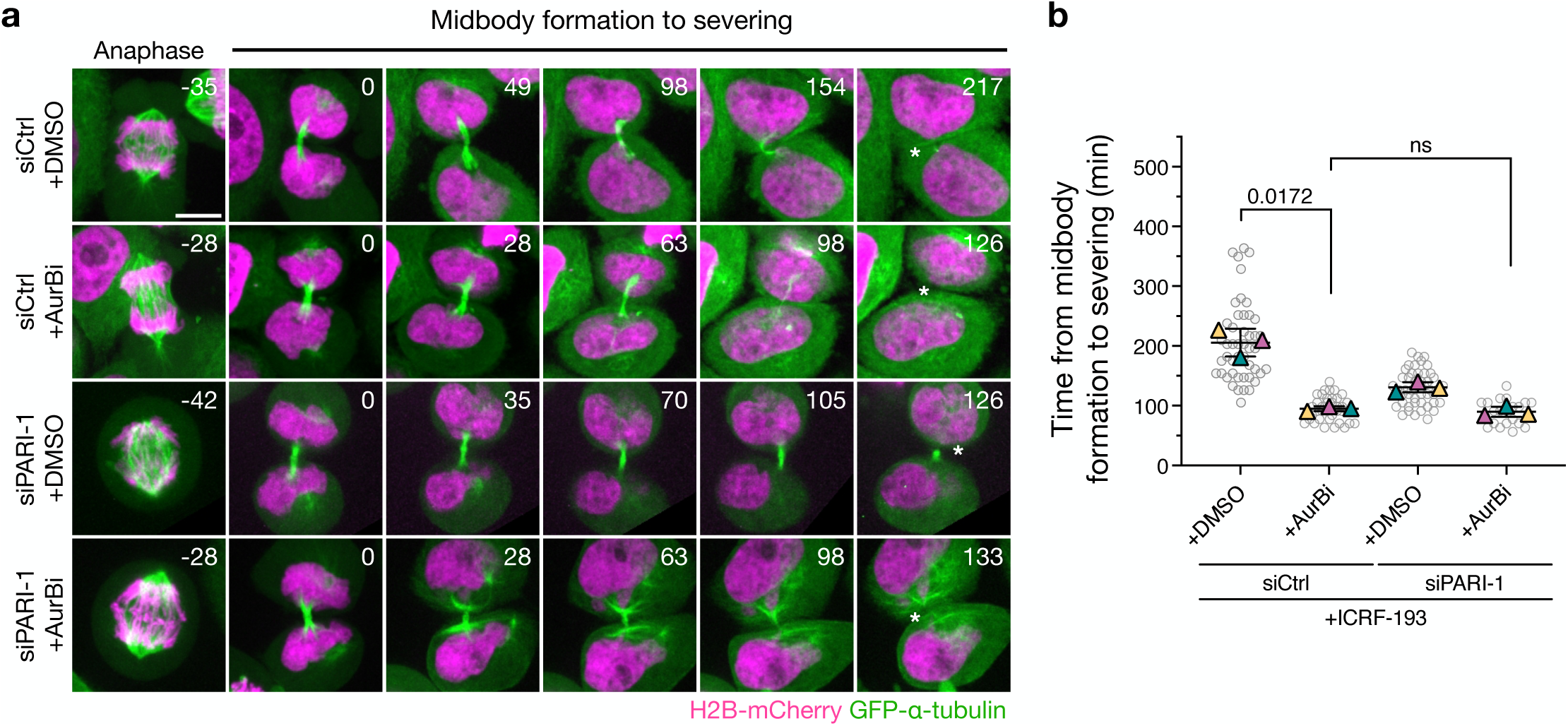
PARI and Aurora-B function in the same pathway to inhibit abscission. (**A**) Cells treated with ICRF-193 progressing from anaphase to midbody formation (GFP-alpha-Tubulin) and its disassembly. The Aurora B inhibitor hesperadin was added 120 min after the start of acquisition when most cells on average had formed the midbody. Asterisk specifies midbody disassembly. Scale bar 5 um. (**B**) Quantification of the midbody lifetime (time from midbody formation to severing). Student’s paired t test (mean ±SD, *P <0.05, **P <0.01; ***P <0.001; ****P <0.0001, n ³20, N =3).

## Discussion

The NoCut checkpoint plays a critical role in ensuring the proper coordination between chromosome segregation and cytokinesis, thus safeguarding genomic integrity during cell division. Disruption of this checkpoint can lead to the damage of chromatin bridges and genomic instability. Understanding the molecular mechanisms behind NoCut is crucial for elucidating how cells protect themselves against such potentially catastrophic events. We hypothesised that DNA helicases might play a role in NoCut signalling because they are well-positioned to mark, either directly or indirectly, chromatin bridges that arise from replication or topological defects, both of which are known triggers of NoCut signalling. Srs2, for example, is recruited to replication forks by SUMOylated PCNA to prevent HR during S phase and G2, and is associated with ultra fine bridges (UFBs). However, its potential functions during mitosis and cytokinesis had not been explored prior to this study.

Our findings reveal that Srs2 plays a crucial role in chromosome segregation during mitosis, particularly following DNA replication stress. Srs2 appears to prevent the accumulation of unreplicated DNA during anaphase, reducing the formation of aberrant DNA structures that could otherwise lead to segregation defects. In the absence of Srs2, cells exhibited increased RPA-coated chromatin bridges associated with delayed bridge resolution. This suggests that Srs2’s activity is important for resolving these aberrant DNA structures caused by replication stress.

Furthermore, Srs2 may play a dual role in protecting cells from DNA damage: first, by promoting the resolution of these structures, and second, by potentially triggering the NoCut checkpoint if these structures remain unresolved by the end of cytokinesis. The fact that *srs2Δ* cells exhibited delayed bridge resolution but did not display the NoCut-dependent abscission delay typically seen in response to chromatin bridges implies that Srs2 might be involved in signalling unresolved structures for NoCut checkpoint activation.

An intriguing aspect of our study is the discovery that Srs2 is required to delay abscission in the presence of both HU-dependent and catenated chromatin bridges. Our observation that depleting the abscission factor Cyk3 reduces DNA damage in *srs2Δ* cells suggests that Srs2’s role in delaying abscission and protecting chromatin bridges from damage is distinct from its anti-recombinogenic function during fork resolution after HU treatment. This indicates that Srs2’s function in cytokinesis (at least in the formation of breaks) is independent of its role in DNA damage repair. Moreover, our experiments with mutants affecting the Srs2-PCNA interaction suggest that Srs2 retention in chromatin bridges may be both necessary and sufficient to enforce an abscission delay. Thus, Srs2 may act as a chromatin-based signal for the presence of chromatin bridges, potentially marking these structures for recognition by the NoCut checkpoint machinery.

In human cells, we found that PARI, the homolog of Srs2, plays a similar role in the abscission checkpoint. Like Srs2, PARI is an inhibitor of HR and is recruited to stalled or collapsed replication forks under replication stress. We also found that PARI is essential for stabilising midbodies and actin structures at the division site, both of which are hallmarks of the NoCut/abscission checkpoint, in the presence of catenated chromatin bridges. This role seems to be specific to the response to chromatin bridges, as it does not participate in the midbody stabilisation seen with nuclear pore complex (NPC) assembly defects, highlighting its specificity within the mammalian NoCut checkpoint.

In summary, our data reveal an evolutionarily conserved function of the DNA helicase Srs2 / PARI in controlling the timing of abscission events in response to chromatin bridges. Despite these insights, the molecular mechanisms by which Srs2 and PARI function in NoCut remain unclear. We do not yet understand how Srs2 promotes the recognition of DNA bridges. One possibility is that its presence on incompletely replicated or catenated DNA at the spindle midzone—where Aurora-B resides—might enhance Aurora-B activity or signalling to downstream NoCut components. While Srs2 has been associated with chromatin bridges, there is no clear evidence that PARI or PCNA localise to these bridges in human cells. Our attempts to detect PARI-GFP or PCNA by immunofluorescence were unsuccessful, although this could be due to their low abundance at these sites.

Interestingly, Topoisomerase II (Topo II) has been suggested to mark catenated bridges for recognition by the human abscission checkpoint (Petsalaki et al., 2023). However, our data indicate that inhibition of Topo II with ICRF-193 remains coupled to key features of the abscission checkpoint, including the stabilization of midbodies and actin structures at the division site. We propose that the low concentration of ICRF-193 used in our study, which is 40-fold lower than in Petsalaki et al., may only partially inactivate Topo II, allowing the formation of chromatin bridges while retaining sufficient Topo II activity to maintain abscission checkpoint function. It is not yet clear whether Srs2/PARI and Topo II mark bridges independently or as part of a larger complex. The possibility of such a complex is supported by yeast two-hybrid studies demonstrating an interaction between Srs2 and Topo II (Chiolo et al., 2005), suggesting that these proteins could collaborate in marking chromatin bridges for NoCut signalling.

Finally, PARI’s association with cancer raises the intriguing possibility that its function in the NoCut checkpoint may contribute to maintaining genomic stability and preventing tumorigenesis. Interestingly, a polymorphism in the human ESCRT-III component CHMP4C, which impairs the abscission checkpoint, has been linked to increased cancer susceptibility (Sadler et al., 2018). Understanding the specific role of PARI in the NoCut checkpoint and how it might intersect with other pathways involved in genomic stability could provide valuable insights into the mechanisms underlying cancer development.

## Acknowledgements

We thank all members of the Mendoza laboratory for their valuable discussions, and Arnaud Echard, Andrés Clemente and Felix Prado for critically reviewing the manuscript. We are grateful to the CRG Advanced Live Microscopy Unit and the IGBMC Imaging Centre, part of the national infrastructure France-BioImaging, supported by the French National Research Agency (ANR-10-INBS-04), as well as the IGBMC Cell Culture and Flow Cytometry Facilities. This work was funded by the European Research Council (ERC) Starting Grant 2010-St-20091118 and grants from the “Ligue Nationale Contre le Cancer” (3FI13531TJQN and 3FI14004UUGH) to M.M., as well as support from the Spanish Ministry of Economy and Competitiveness, ‘Centro de Excelencia Severo Ochoa 2013–2017’, SEV-2012-0208 to the CRG. MD was supported by a LabEx PhD fellowship from IGBMC and a fellowship from Ligue Nationale Contre le Cancer. NB was supported by a Marie Skłodowska-Curie postdoctoral fellowship. Additionally, this work, part of the Interdisciplinary Thematic Institute IMCBio+ 2021-2028 program of the University of Strasbourg, CNRS, and Inserm, was supported by IdEx Unistra (ANR-10-IDEX-0002), SFRI-STRAT’US project (ANR-20-SFRI-0012), and EUR IMCBio (ANR-17-EURE-0023) under the France 2030 Program framework.

## Author Contributions (CRediT author statement)

**MD**: Conceptualization, Data Curation, Investigation, Methodology, Validation, Visualization, Writing – Original Draft. **NB**: Conceptualization, Data Curation, Investigation, Methodology, Validation, Visualization. **AF**: Investigation, Methodology. **CS**: Investigation, Methodology, Formal Analysis, Visualization. **MM**: Conceptualization, Data Curation, Funding Acquisition, Supervision, Visualization, Writing – Original Draft, Writing – Review & Editing.

## Materials and Methods

### Yeast strains and culture

All *Saccharomyces cerevisiae* strains are derived from S288c (**Table 1**). Gene deletions and insertions were generated by standard PCR methods (Janke et al. 2004) or through crossing. Cells were grown in YPDA media (yeast extract, peptone, 2% glucose, and adenine) at the permissive temperature of 30°C, and temperature-sensitive strains were grown at 25°C and shifted to 37°C to inactivate the protein. Dicentric strains were grown in YPDA supplemented with 2% galactose instead of glucose.

For all synchronizations, cells were grown overnight in YPDA at the permissive temperature, diluted to OD_600_ =0.1, and grown for 3 hours to reach the log phase. To induce expression and localization of CAAX-GFP to the plasma membrane, 90 nM β-estradiol was added to the media for 3 hours. To generate replication stress 200 mM HU was added to the media for 3 hours. To arrest cells in G1, cells synchronised with 20 μm/ml α-factor for 2 hours. To release from the G1 block or HU-treatment cells were washed twice in freshly prepared and pre-heated minimal synthetic (Yeast nitrogen base, 2% glucose, essential amino acids) media and immediately plated on concanavalin A-coated Lab-Tek chambers for microscopy. For analysis of Rfa2-GFP during cell division, visual inspection showed that all cells displayed GFP-labeled nuclear foci during interphase, as described (Ivanova et al., 2020). Cells that retained foci after the onset of nuclear elongation were classified as “chromosome segregation with RPA foci”.

### Human cell lines and culture conditions

All cell lines were cultured at 37°C in a 5% CO2 humidified incubator. Standard media, DMEM GlutaMAX (4.5 g/L glucose) supplemented with 10% foetal calf serum, 1% penicillin, and streptomycin was used to culture all cell lines. The HeLa Kyoto cell line was obtained from the in-house cell culture facility. HeLa stable cell lines were kindly shared by Dr. Daniel Gerlich and were grown in standard media with additional selective antibiotics.

### Plasmid and siRNA transfection

A FLAG-PARI Gateway Destination plasmid was kindly gifted by Dr. Peter Burkovics, and the sequence of PARI was further cloned into pcDNA3.1(+) plasmid by the molecular biology facility at IGBMC. All plasmid cloning and primer design were performed by the molecular biology facility. An eGFP fluorophore was introduced at the N-terminal of PARI. For the siPARI-1 resistant version of eGFP-PARI three silent mutations were introduced at the seed region of the siPARI-1 target site.

To generate stable cell lines, HeLa Kyoto cells were transfected with linearized pcDNA3.1(+) derived plasmids using X-tremeGENE 9 DNA Transfection Reagent (ROCHE) according to the manufacturer’s protocol. Transfected cells were selected for 2-3 weeks in standard media supplemented with G418 (0.8 mg/mL). Transgene-positive cells were isolated by FACS (FACS ARIA, BD Biosciences) or by manually isolating single colonies. Expression was validated by PCR, IF, and WB.

All siRNAs used are of the siGENOME product line from Dharmacon and were purchased from Horizon (**Table 2**). To knockdown PARI, resuspended cells were (reverse) transfected using 25 nM siRNA and Lipofectamine RNAiMAX (Invitrogen) according to the manufacturer’s protocol, the siRNA-lipid complexes were removed after 7 hours, and a second (forward) transfection was performed 24 hours after the first one. For co-depletion of PARI and NUP153, 10 nM siNUP153 was used together with 25 nM siCtrl or siPARI-1 24 hours after reverse transfection with siCtrl and siPARI-1, and the amount of Lipofectamine RNAiMAX was adjusted accordingly.

### Quantitative PCR and analysis

RNA was extracted using RNAeasy kit (Qiagen), and 2.5 µg RNA was incubated with ezDNase for 10 min at 37°C before performing reverse transcription using superscript IV VILO Master Mix according to the manufacturer’s protocol. Alternatively, DNase I was used, and reverse transcription was performed with random hexamer and oligo(dT) primers, SuperScript IV, and RNaseOUT Recombinant RNase Inhibitor according to the manufacturer’s protocol. SYBR Green-based qPCR and analysis were carried out on the LightCycler480.

All primer pairs were validated with PCR and serial dilution to determine the primer efficiency, only those that showed >90% amplification efficiency were used for analysis (**Table 3**). The Ct value was obtained from the LighCycler480 software and used to measure the relative expression (RE) of each sample, which was calculated using the following equation

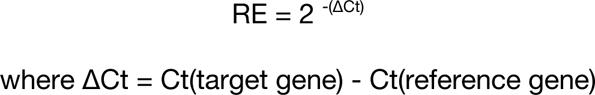

The RE of the test sample is then normalized to the RE of the control.

### Immunofluorescence microscopy and image analysis

20.000 cells were seeded on 13 mm sterile glass coverslips in 24-well culture plates and fixed with 4% PFA for 15 min at RT. PFA was washed out three times with 1x PBS and cells were permeabilized in 0.2% Triton X-100 in 1x PBS, for 10 min at RT, followed by a 1-hour block with 3% BSA in 1x PBS at RT with gentle agitation. After blocking, cells are incubated with primary antibodies (**Table 4**) for 1 hour at RT or overnight at 4°C, followed by three washes with 1x PBS. Cells are then subjected to secondary antibodies (**Table 4**) and DAPI for 1 hour at RT, followed by three washes with 1x PBS. Excess 1x PBS was carefully tapped off from the coverslips and mounted on a glass slide with Vectashield mounting medium (Vecta Laboratories) or ProLong Gold (Invitrogen). The HCX PL APO 63x/1.40 OIL PH3 C3 objective was used on an upright motorised Leica DM 4000 B equipped with CoolSNAP HQ2 camera.

### Time-lapse fluorescence spinning disk confocal microscopy

For abscission assays, HeLa cells were seeded 3 days in advance on a 4-chamber 35 mm glass bottom dish (Cellvis) and placed in a preheated chamber at 37°C and 5% CO2 for time-lapse microscopy. For cell lines that do not stably express fluorescently tagged Tubulin, cells were incubated with 15 nM SiR-Tubulin (Spirochrome) and 1 μM Verapamil for 9 hours in media before acquisition. All time-lapse acquisitions were set up using MetaMorph software. Images were acquired using 63x water objective (HC PL APO 63x/1,20 W CORR CS2) with Leica water immersion micro dispenser connected to a Bartek extended micropump on an inverted Leica DMI8 microscope equipped with Yokogawa CSU W1 spinning disk and Evole 512 camera or an inverted Nikon eclipse equipped with Yokogawa CSU X1 spinning disk and photometric prime 95B camera. Z-stack (15 μm range, 0.3 μm step size) images were acquired every 5-7 min for 14 hours. For time-lapse microscopy of Aurora B inhibition with hesperadin, media with a final concentration of 100 nM hesperadin was added on top of the cells in between intervals.

### Correlative light electron microscopy

Adherent HeLa Kyoto stably expressing H2B-mCherry GFP-α-Tubulin cells were first cultured on laser micro-patterned Aclar^®^ supports (Spiegelhalter et al., 2014), synchronised with double thymidine and released for 12 hours. Cells were then fixed with 1% glutaraldehyde and 4% formaldehyde in a 0.1M phosphate buffer for 30 minutes. Cells in cytokinesis with or without a catenated chromatin bridge were selected, precisely located, and imaged by an inverted Nikon eclipse equipped with a photometric prime 95B camera. At the end of the experiment, cells were immediately fixed with 2.5% glutaraldehyde and 4% formaldehyde in 0.1 M phosphate buffer for 1 hour (or longer) at 4°C, rinsed in buffer, and followed by 1-hour postfixation in 1% osmium tetroxide [OsO_4_] reduced by 0.4% potassium hexacyanoferrate (III) [K_3_Fe(CN)_6_] in H2O at 4°C. Samples were rinsed in distilled water and stained with 1% tannic acid for 30 minutes on ice and after extensive rinses, with 2% uranyl acetate for 1 hour at room temperature, rinsed in water. Samples were dehydrated with increasing concentrations of ethanol (25%, 50%, 70%, 90%, and 3×100%), and embedded with a graded series of epoxy resin. Samples were finally polymerized at 60°C for 48 hours. Ultrathin serial sections (70nm) were picked up on 1% pioloform coated copper slot grids and observed with a Philips CM12 operated at 80kV equipped with an Orius 1000 CDD camera (Gatan, Pleasanton, USA).

### Statistical methods

Graphpad Prism software was used to generate graphs and perform statistical tests. Mann-Whitney was used on datasets that did not follow a normal distribution. For experiments with at least three independent biological replicates, Students paired t-test was used. Fisher’s exact test was performed on data that was pooled from at least two biological replicates to compare fractions.

