## Supplementary Figures and Tables for "Srs2/PARI DNA helicase mediates abscission inhibition in response to chromatin bridges in yeast and human cells"

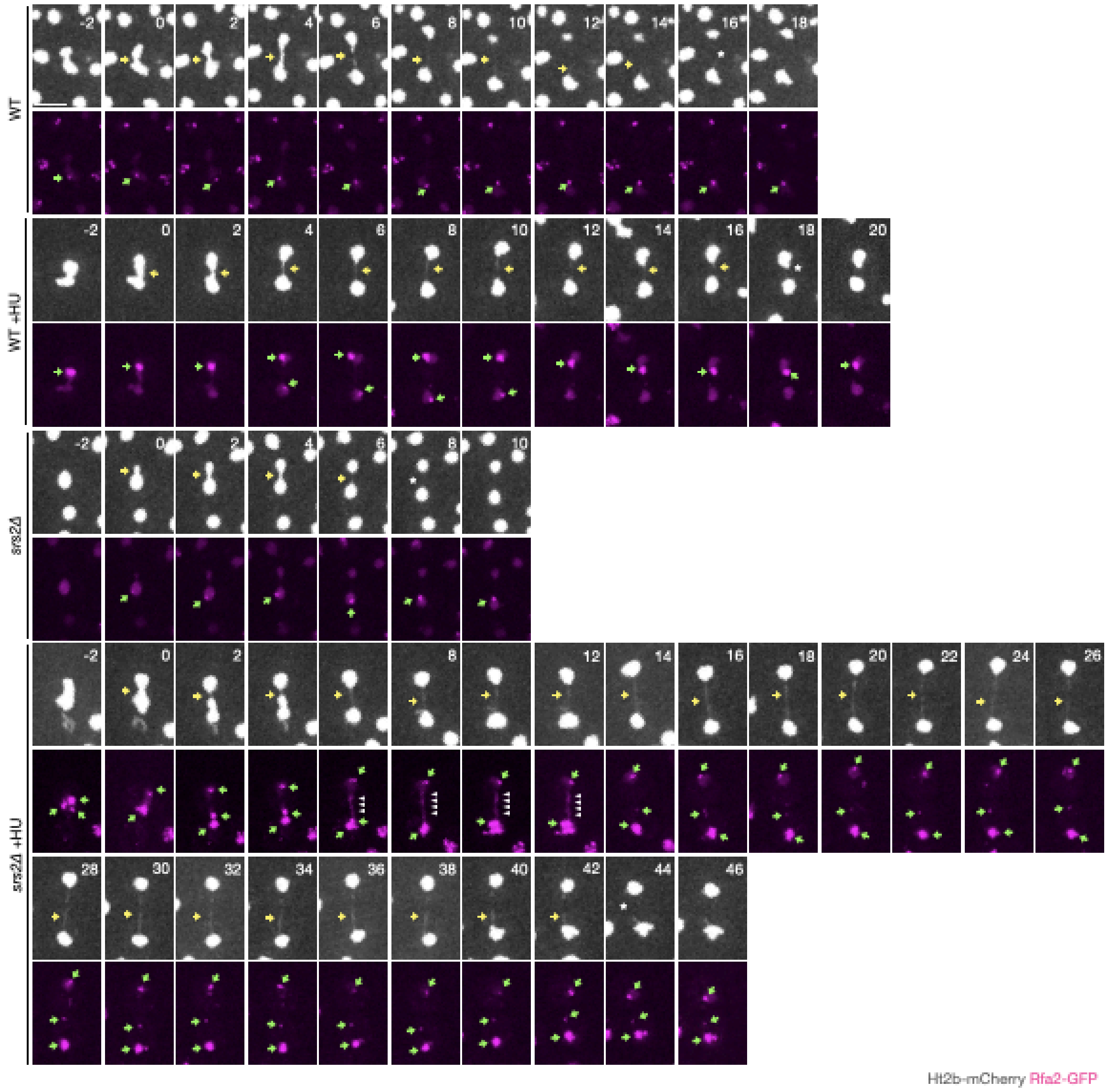


**Supplementary Figure 1.** Chromosome segregation (Htb2-mCherry) and RPA foci formation (Rfa2-GFP) in cells of the indicated strains, shown in Figure 1A, including all time points, with and without previous exposure to HU. Only cells with RPA foci are shown. Yellow arrows indicate chromatin bridges, green arrows point towards RPA foci, white arrowheads mark RPA bridges and asterisks note bridge resolution. Scale bar: 5 microns.

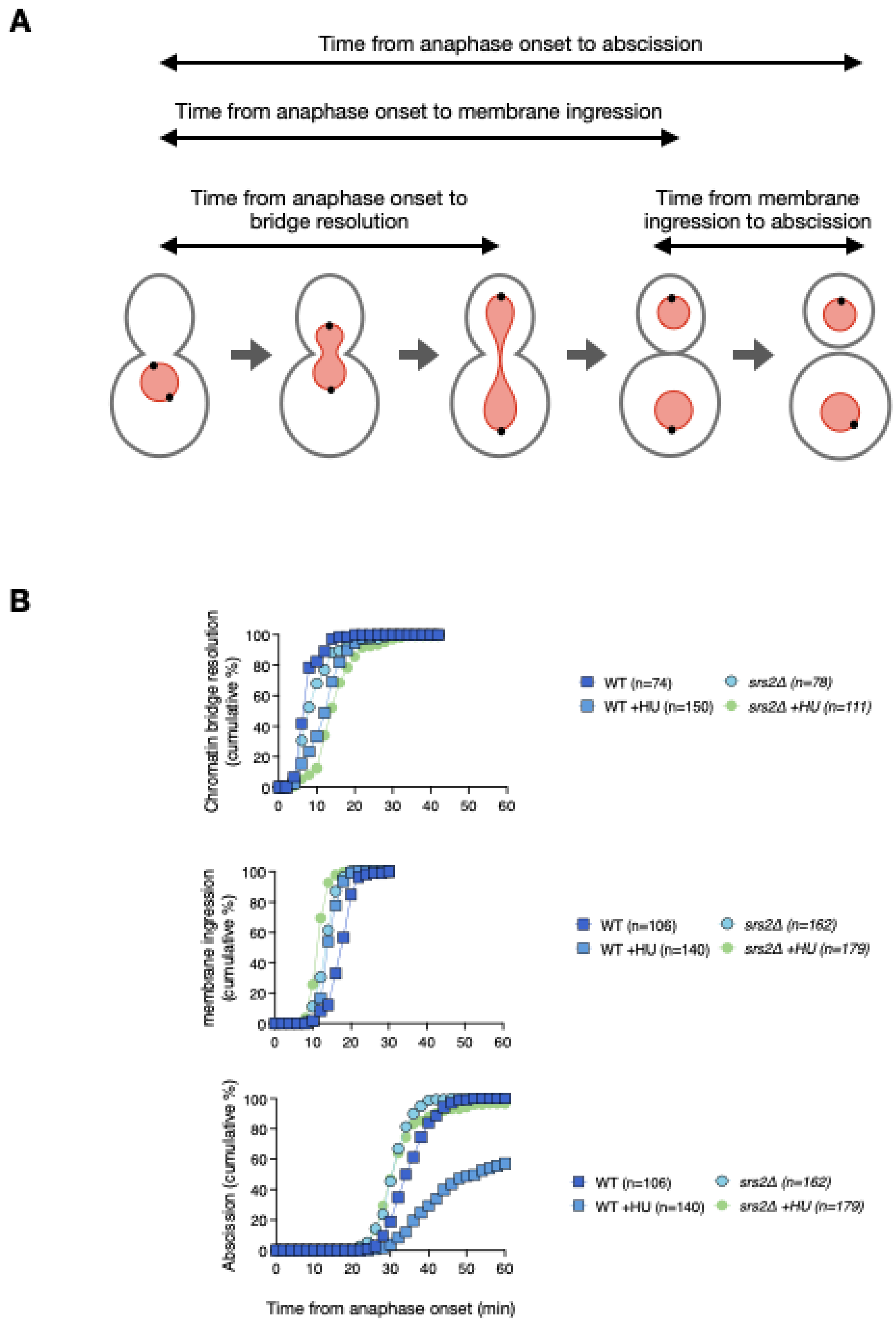


**Supplementary Figure 2. Timing of chromatin bridge resolution, membrane ingression, and abscission in wild-type and srs2Δ cells.** (**A**) Schematic representation of key cell division events, including chromatin bridge resolution, membrane ingression, and abscission. The time intervals measured in the analysis are indicated by horizontal arrows. The nucleus is in red, the spindle pole bodies are black circles, and the plasma membrane is in grey. (**B**) Cumulative frequency plots showing the timing of chromatin bridge resolution (top), membrane ingression (middle), and abscission (bottom) in wild-type (WT) and *srs2Δ* cells, with or without hydroxyurea (HU) treatment. The number of cells analyzed for each condition is indicated in the legend.

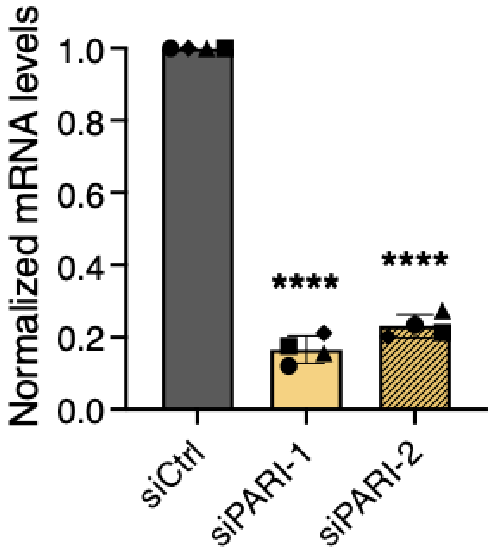


**Supplementary Figure 3.** RT-qPCR measures the mRNA levels of PARI in HeLa cells after transfecting twice with 25 nM siCtrl, siPARI-1 and siPARI-2. Relative mRNA levels have been normalised to siCtrl. One sample t and Wilcoxon test (mean ±SD, *P <0.05, **P <0.01; ***P <0.001; ****P <0.0001, N =4).

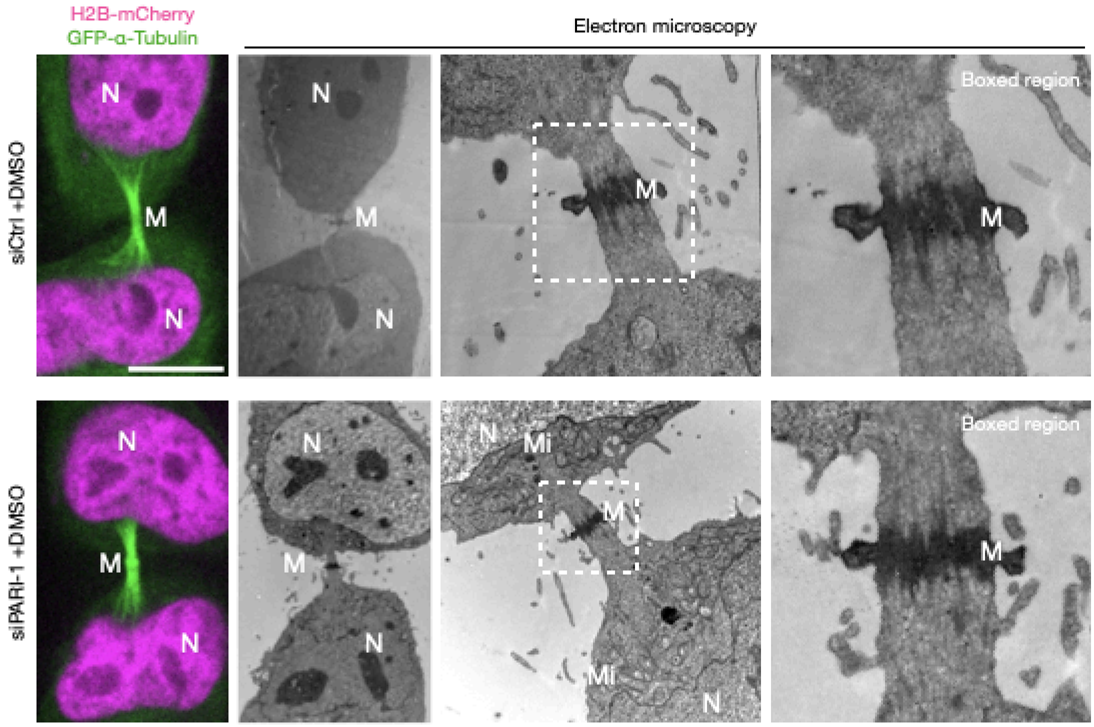


**Supplementary Figure 4.** Cells stably expressing H2B-mCherry and GFP-alpha-Tubulin, transfected with control or PARI-1-specific siRNAs. Boxed regions zoom in on the Flemming body. N = nucleus, M = midbody, Mi = mitochondria.

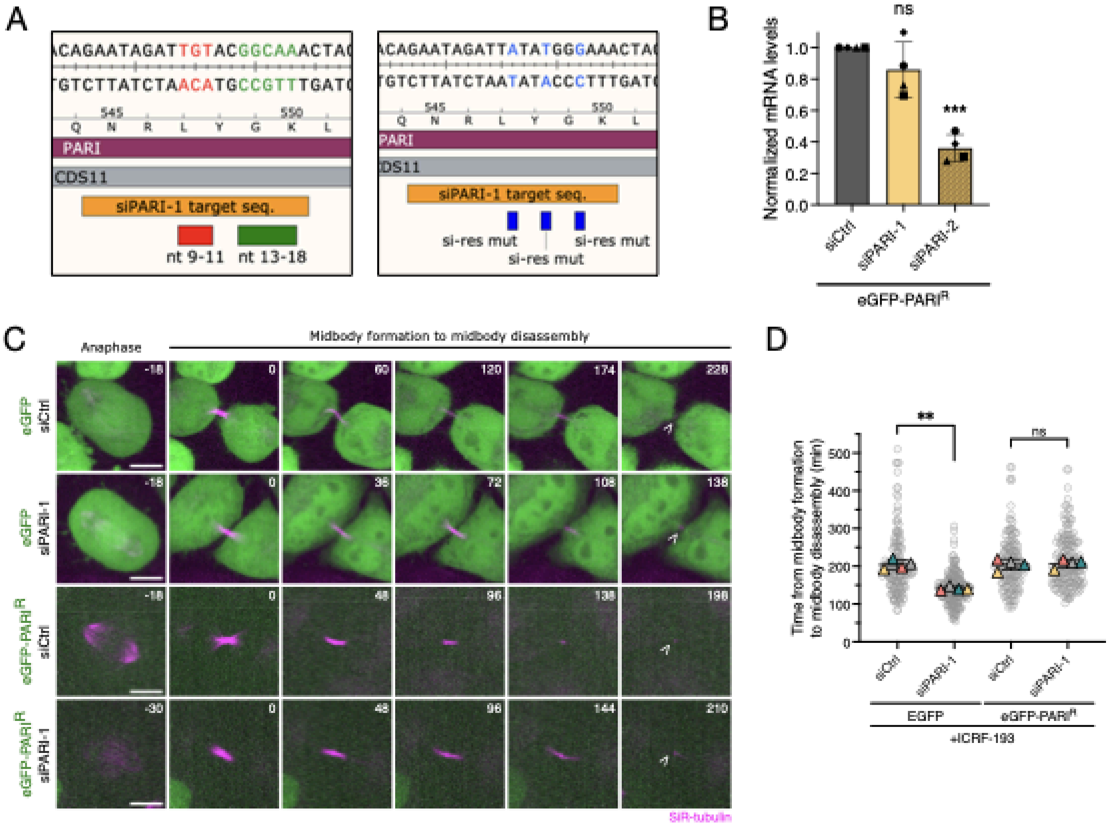


**Supplementary Figure 5.** (**A**) Left panel: WT siPARI-1 target sequence, features highlighting seed sequences. Right panel: siPARI-1 resistant PARI sequence, features show where three silent point mutations were introduced by cloning. (B) RT-qPCR measures the mRNA levels of PARI in HeLa cells after transfecting twice with 25 nM siCtrl, siPARI-1 and siPARI-2. Relative mRNA levels have been normalised to siCtrl. One sample t and Wilcoxon test (mean ±SD, *P <0.05, **P <0.01; ***P <0.001; ****P <0.0001, N =4). (**C**) Cells treated with ICRF-193 progressing from anaphase to midbody formation (SiR-Tubulin) and its disassembly. Cells have been synchronised and treated as illustrated in Figure 5. Open arrowhead specifies midbody disassembly. Scale bar 5 mm. (**D**) SuperPlot showing the quantification of the midbody lifetime (time from midbody formation to disassembly). Student's paired t test (mean ±SD, *P <0.05, **P <0.01; ***P <0.001; ****P <0.0001, n =165, N =4).

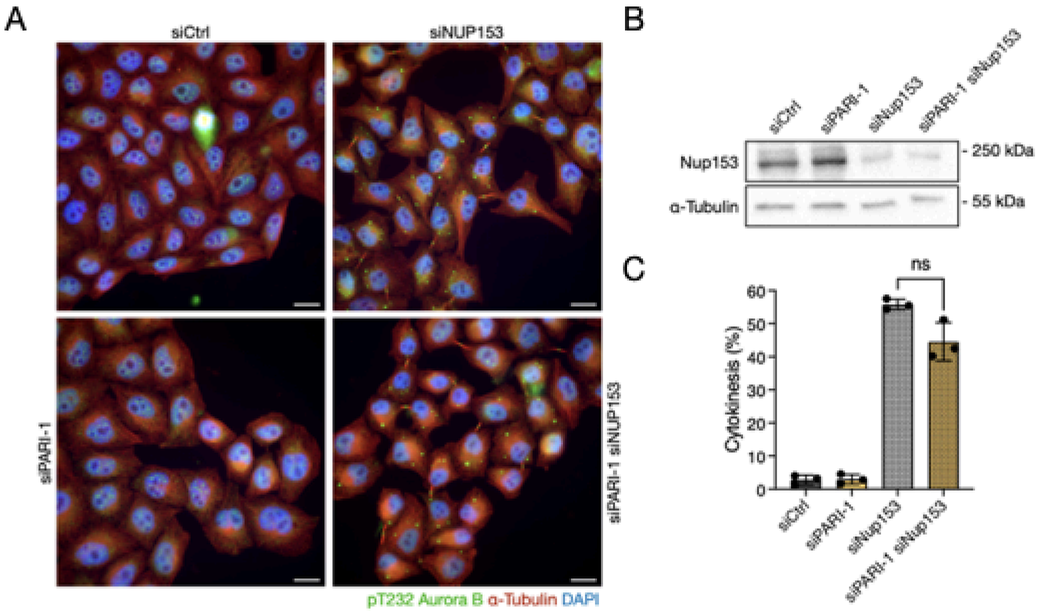


**Supplementary Figure 6.** (**A**) Cells transfected with siCtrl, siNUP153, siPARI-1 and a combination of siPARI-1 + siNUP153. Cells were synchronised with thymidine for 24 hours, released for 16 hours and fixed. (**B**) Western blot showing partial depletion of NUP153. Membrane probed with a NUP153 antibody and alpha-Tubulin as a loading control. (**C**) Quantification of the fraction of cells arrested in cytokinesis. Student's paired t test (mean ±SD, *P <0.05, **P <0.01; ***P <0.001; ****P <0.0001, n $^{3}$789, N =3).

**Table 1: *Saccharomyces cerevisiae* strains**

| YMM ID | Background | Genotype |
|---|---|---|
| 1335 | S388C | ADEGV:URA3<br>Gal1:GFP-CAAX:HIS3<br>SPC42-GFP:HphMX<br>leu2 lys2-801 ade2-101 trp1Δ63 |
| 3399 | S388C | SRS2::NAT<br>ADEGV:URA3<br>Gal1:GFP-CAAX:HIS3<br>SPC42-GFP:HphMX<br>leu2 lys2-801 ade2-101 trp1Δ63 |
| 2378 | S388C | top2-4<br>ADEGV:URA3<br>Gal1:GFP-CAAX:HIS3<br>SPC42-GFP:HphMX<br>leu2 lys2-801 ade2-101 |
| 3401 | S388C | SRS2::NAT<br>top2-4<br>ADEGV:URA3<br>Gal1:GFP-CAAX:HIS3<br>SPC42-GFP:HphMX<br>leu2 lys2-801 ade2-101 |
| 3949 | BY4741 | RFA2-GFP:HIS<br>Htb2-mCherry:HphMX |
| 6161 | BY4741 | SRS2::NAT<br>RFA2-GFP:HIS<br>Htb2-mCherry:HphMX |
| 4581 | S288C | POL30-S115P<br>ADEGV:URA3<br>pGal1:GFP-CAAX:HIS3<br>SPC42-GFP:HphMX<br>leu2 lys2-801 ade2-101 trp1Δ63 |

| | | |
|---|---|---|
| 3298 | S288C | SRS2ΔSIM::NAT<br>top2-4<br>ADEGV:URA3<br>pGal1:GFP-CAAX:HIS3<br>SPC42-GFP:HphMX<br>leu2 lys2-801 ade2-101 |
| 3960 | S288C | SRS2ΔSIMΔPIP::NAT<br>top2-4<br>ADEGV:URA3<br>pGal1:GFP-CAAX:HIS3<br>SPC42-GFP:HphMX<br>leu2 lys2-801 ade2-101 |
| 2833 | Mostly S228C mixed with W303 | ChrXII(1059):nat:ChrIV(19.5)<br>pGal:CEN4:KanMX4<br>Htb2-mCherry:URA<br>pGal1:GFP-CAAX:TRP1<br>ura3-52 his3Δ200 leu2 lys2-801 ade2-101 trp1Δ63 |
| 4578 | Mostly S228C mixed with W303 | ELG1::HYG<br>ChrXII(1059):nat:ChrIV(19.5)<br>pGal:CEN4:KanMX4<br>Htb2-mCherry:URA<br>pGal1:GFP-CAAX:TRP1<br>ura3-52 his3Δ200 leu2 lys2-801 ade2-101 trp1Δ63 |

**Table 2. siRNA sequences for gene knockdown**

| siRNA | Sequence 5'-3' |
|---|---|
| siCtrl pool | UAAGGCUAUGA AGAGAUAC,<br>AUGUAUUGGCCU-GUAUUAG,<br>AUGAACGUGAAUUGCUCAA,<br>UGGUUUACAUGUCGA-CUAA |
| siPARI-1 | GAATAGATTGTACGGCAAA |
| siPARI-2 | CCAAGGACAAGTTGATTTC |
| siNUP153 | GAGGAGAGCUCUAAUAUUA |

**Table 3. Primer sequences for qPCR**

| Target gene | Forward/Reverse | Sequence 5'-3' |
|---|---|---|
| ACTIN B | Forward | AGGCACCAGGGCGTGAT |
| ACTIN B | Reverse | GCCCACATAGGAATCCTTCTGAC |
| PARI | Forward | GCATCAAAGCCTTTGTGTG |
| PARI | Reverse | CCTGTCTGACTGGTTGAT |

**Table 4. Antibodies used for western blot and immunofluorescence**

| Primary antibodies | | | |
|---|---|---|---|
| Raised in | Against | Reference | Dilution |
| Mouse | a-Tubulin | T9026 | 1:1000 (IF, PFA) |
| Mouse | Nup153 | ab24700 | 1:1,000 (WB) |
| Rabbit | a-Tubulin | ab52866 | 1:1000 (IF, PFA)<br>1:5,000 (WB) |
| Secondary antibodies | | | |
| Goat | Anti-mouse IgG Alexa Fluor Plus 488 | A32723 | 1:500 |
| Goat | Anti-mouse IgG Alexa Fluor 568 | A-11031 | 1:500 |

| | | | |
|---|---|---|---|
| **Goat** | **Anti-rabbit IgG Alexa Fluor 488** | **A32731** | **1:500** |
| **Goat** | **Anti-rabbit IgG Alexa Fluor 647** | **A32733** | **1:500** |
| **Goat** | **Anti-mouse IgG HRP** | **170-6516** | **1:10,000** |
| **Goat** | **Anti-rabbit IgG HRP** | **31460** | **1:10,000** |
